# What makes human cortical pyramidal neurons functionally complex

**DOI:** 10.1101/2024.12.17.628883

**Authors:** Ido Aizenbud, Daniela Yoeli, David Beniaguev, Christiaan PJ de Kock, Michael London, Idan Segev

**Affiliations:** The Edmond and Lily Safra center for Brain Sciences (ELSC), The Hebrew University of Jerusalem, Jerusalem, Israel; Department of Integrative Neurophysiology, Center for Neurogenomics and Cognitive Research (CNCR), Neuroscience Campus Amsterdam, VU Amsterdam; Department of Neurobiology, The Hebrew University of Jerusalem, Jerusalem, Israel

**Author notes:** Denotes equal contributions (Ido Aizenbud, Daniela Yoeli).

**Keywords:** single neuron computation, dendritic computation, functional complexity, human neurons, rat neurons, cortical pyramidal neurons, compartmental modeling, biophysical modeling, deep neural networks

## Abstract

Humans exhibit unique cognitive abilities within the animal kingdom, but the neural mechanisms driving these advanced capabilities remain poorly understood. Human cortical neurons differ from those of other species, such as rodents, in both their morphological and physiological characteristics. Could the distinct properties of human cortical neurons help explain the superior cognitive capabilities of humans? Understanding this relationship requires a metric to quantify how neuronal properties contribute to the functional complexity of single neurons, yet no such standardized measure currently exists. Here, we propose the Functional Complexity Index (FCI), a generalized, deep learning-based framework to assess the input-output complexity of neurons. By comparing the FCI of cortical pyramidal neurons from different layers in rats and humans, we identified key morpho-electrical factors that underlie functional complexity. Human cortical pyramidal neurons were found to be significantly more functionally complex than their rat counterparts, primarily due to differences in dendritic membrane area and branching pattern, as well as density and nonlinearity of NMDA-mediated synaptic receptors. These findings reveal the structural-biophysical basis for the enhanced functional properties of human neurons.

## Introduction

It is generally accepted that the unique cognitive capabilities of humans arise from a combination of many attributes. At the macroscale, these attributes might include the large number of computational elements (neurons/glial cells), the intense region-to-region connectivity and the human regional specialization (Gabi et al., 2016; Axer and Amunts, 2022, Rockland, 2023). At the microscale, it was suggested that human specific transcriptomic features contribute to these capabilities (Jostard et al., 2023). It was also argued that human cognition might be supported by the evolution of new cell types (Berg et al., 2021) and the unique morphological and biophysical properties of human cortical neurons (Galakhova et al., 2022). Indeed, studies have identified numerous distinctive properties in human cortical neurons (Spruston, 2008; DeFelipe, 2011; Mohan et al., 2015; Deitcher et al., 2017; Eyal et al., 2018; Mihaljevic et al., 2021; Galakhova et al., 2022; Han et al., 2023; Hunt et al., 2023). However, the impact of this cellular-level complexity on the computational capabilities of the neuron, and consequently on the entire neuronal system, remains unclear.

Already Ramon y Cajal noticed that human cortical neurons are particularly large and morphologically complex (Ramón y Cajal et al., 1988). Over the past two decades, numerous studies have systematically compared the dendritic geometry of human cortical and hippocampal neurons with that of other species, particularly rodents. Human cortical neurons are generally characterized by large dendritic trees with elongated branches, especially the terminal branches of the basal dendrites (Deitcher et al., 2017), and extensive arborization (Spruston, 2008; DeFelipe, 2011; Mohan et al., 2015; Eyal et al., 2018; Mihaljevic et al., 2021; Galakhova et al., 2022; Han et al., 2023; Hunt et al., 2023; Oláh et al., 2024). The large and extensive dendritic arborization provides a large surface area for receiving and processing synaptic inputs, and supports sampling from a diverse array of inputs. Furthermore, Large dendritic extensions lead to electrical decoupling between dendritic regions that give rise to dendritic compartmentalization, which allows distinct regions of the dendritic tree to operate as semi-independent computational subunits (Polsky et al., 2004; Beualieu-Laroche et al., 2018; Eyal et al., 2018; Beualieu-Laroche et al., 2021; Otor et al., 2022).

In addition to the morphological distinctions between cortical neurons in rats and humans, several biophysical and synaptic attributes differ across species. Specific membrane properties are one such attribute (Eyal et al., 2016; Eyal et al., 2018; Chameh et al., 2023); other attributes include the time-dependent dynamics of the synaptic connection (Mansvelder et al., 2019; Testa-Silva et al., 2010) and nonlinear dendritic properties (Gidon et al., 2020), particularly the density and steepness of the voltage-dependence of N-methyl-D-aspartate (NMDA) receptors - both were found to be larger in human cortical pyramidal neurons compared to rodents (Eyal et al., 2018; Hunt et al., 2023; but see Testa-Silva et al., 2022). These biophysical properties of human dendrites are likely to enhance their computational capabilities, e.g., by increasing the number of independent nonlinear dendritic functional subunits (Mel, 1992; Schiller et al., 2000; Poirazi and Mel, 2001; Poirazi et al., 2003a; Poirazi et al., 2003b; Polsky et al., 2004; London and Hausser, 2005; Branco et al., 2010; Eyal et al., 2018; Leleo and Segev, 2021; Tang et al., 2023).

What is critically missing to advance the understanding of how various neuronal characteristics contribute to the functional capabilities of the neuron is a systematic measure that quantifies the functional complexity of neurons, particularly human neurons. Several approaches have been used to systematically assess the computational complexity of single neurons. Poirazi and Mel (2001) used simplified conceptual neuron models to show that both the increased nonlinearity of dendritic integration and the sheer number of bifurcation branches increase a neuron’s memory capacity. Eyal et al. (2018) showed, using detailed compartmental models, that human L2/3 cortical neurons indeed have a larger number of independent nonlinear dendritic subunits compared to rodents. Ujfalussy et al. (2018) captured dendritic computations under *in vivo*-like conditions using models of increasing complexity and used them to characterize input integration of several neuronal types, though only considering the subthreshold activity of the neuron. Recently, Beniaguev et al. (2021) used a deep neural network (DNN) model analogue of a rodent’s L5 cortical neuron to assess the I/O complexity of this neuron, demonstrating the critical role of NMDA-dependent synapses in determining how deep the analogue DNN is. However, a systematic and quantitative exploration of the influence of the full morphological and biophysical range of the neuron’s properties on its I/O computational complexity is not yet available.

To address this gap, we employed a modern machine-learning approach based on Beniaguev et al. (2021). We introduce the functional complexity index (FCI), a novel metric for assessing the functional complexity of neurons. The FCI allows to extract the factors contributing to a neuron’s computational complexity and enables comparisons of I/O complexity across different neuronal types. This comparative analysis offers new insights into fundamental differences in the computational capabilities of cortical neurons between humans and rats, as well as among neurons in different cortical layers, shedding light on the relationship between neurons’ morpho-electrical features and their functional complexity.

## Results

Figure 1 summarizes the steps towards defining the complexity of a given biophysically detailed model of a neuron. First, we generated an I/O dataset for the respective biophysical neuron model by driving it with a large set of synaptic inputs over all of its dendritic tree (Figure 1A, B) and collecting both the subthreshold and suprathreshold voltage output at the soma (Figure 1C, black trace, and see (Beniaguev et al., 2021)). Next, we constructed a fixed, three-layer temporally convolutional neural network (TCN, Bai et al., 2018) with 128 neurons per hidden layer (Figure 1B, and see **Methods**) and trained it to approximate the output of the biophysical neuron model for the same synaptic inputs (Figure 1C, blue trace and see **Methods**). As apparent in Figure 1C, some of the spikes produced by the biophysical model were captured by the respective DNN, while others were missed. The overall quality of the performance of the TCN is assessed by the Area Under Curve (AUC) of the Receiver Operator Characteristic (ROC) curve of spike prediction, with 1 ms temporal resolution (see **Methods**). The more complex the neuron model is, the more spikes are missed by the respective DNN, and the smaller the AUC is. Namely, the more complex the I/O of the neuron, the less accurate the selected fixed DNN is in replicating its I/O properties.

**Figure 1.**
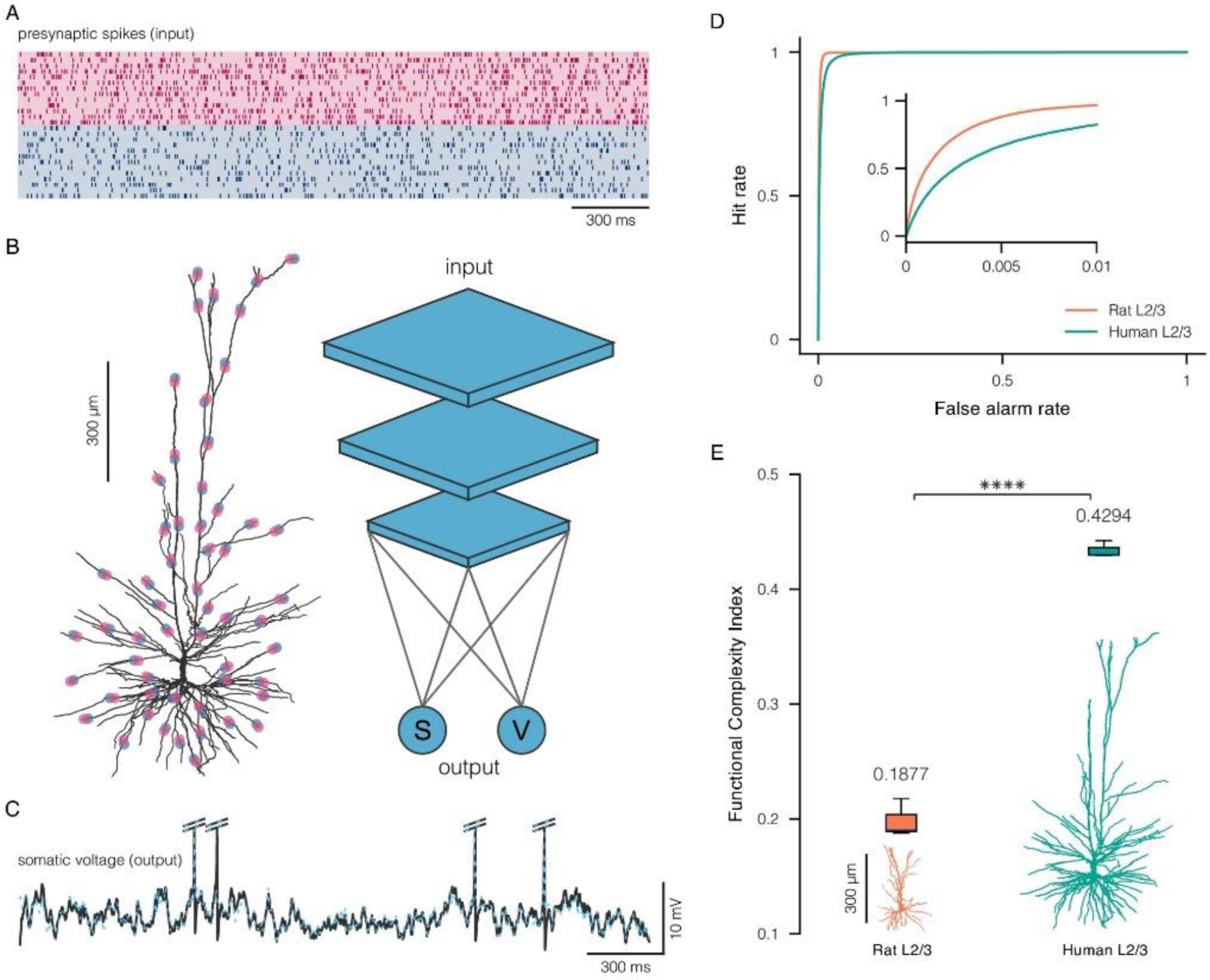
Steps in quantifying the functional complexity of neurons. **A.** Raster plot of random input spikes activating excitatory (red) and inhibitory (blue) synapses distributed over the dendritic tree of the modeled neuron. **B.** Exemplar human layer 2/3 pyramidal neuron (left) and schematics of a three-layer temporal convolutional network *(TCN,* right) that is trained to replicate as closely as possible both the subthreshold (**V**) and spiking activity (**S**) of the biophysical model of the cell shown on the left. **C.** Voltage output (black) of the biophysical model of human L2/3 neuron shown above, and the output of the respective TCN (blue). **D.** Receiver operator characteristic (ROC) curve of spike prediction by a fixed, three-layer TCN (see **Methods**) of the exemplar human layer 2/3 neuron shown in B (green) and of an exemplar rat layer 2/3 neuron shown in D (orange). The area under each of these (green and orange) curves (AUC) indicates the prediction accuracy of the TCN at 1 ms precision, the larger the AUC the better the prediction. **E.** Functional complexity index (FCI) of the exemplar L2/3 human (green) and rat (orange) cortical pyramidal neurons. The FCI ranges from 0 to 1, where 1 is the most complex neuron. **** p value smaller than 0.0001.

Figure 1D shows the result for two exemplar modeled cells: L2/3 cortical pyramidal neurons from human and rat brains (see Figure 1E, bottom, rat in orange and human in green). These two biophysical models had identical passive dendritic properties. The synaptic parameters for these two models respectively match experimental data from human and rat (see **Methods**). For each neuron model, we repeated the training and testing processes of the respective DNN three times, with three different random initial conditions (see **Methods**).

The AUC is inversely related, in a nonlinear manner, to the complexity of the neurons’ I/O properties. The more complex the I/O is, the smaller the respective AUC (where AUC = 1 corresponds to a perfect fit and lowest complexity). To obtain a measure that monotonically increases with I/O complexity, we defined the Functional Complexity Index (FCI) as a monotonically decreasing function of the AUC:

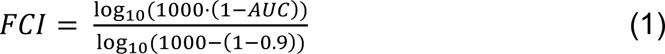

The FCI increases with complexity, and it is approximately linear in the relevant regime; it assumes values close to 0 when the AUC is close to 0.999 (an excellent prediction performance for biophysical neuron models), and values close to 1 when the AUC is close to 0.9 (which indicates poor prediction performance for biophysical neuron models (see **Methods**). For the two exemplar neurons shown in Figure 1D, the FCI is significantly larger for the human L2/3 pyramidal neuron in comparison with the rat L2/3 pyramidal neuron (0.4294 vs 0.1877, two-sided t-test p=2.221e-05).

We next computed the FCI for 24 neuron models: 12 rat pyramidal neurons and 12 human pyramidal neurons spanning all six cortical layers (Figure 2). We used three exemplar cells for each cortical layer (layer 2/3, layer 4, layer 5 and layer 6). In these simulations, all biophysical models have identical passive dendritic properties, but the synaptic models were different for humans versus rats (see **Methods**). The modeled neurons are presented in Figure 2A, bottom, along with their respective FCI (top). Human pyramidal neurons attain much higher complexity levels than rat pyramidal neurons (Figure 2C). The average FCI of all 12 human and 12 rat neurons modeled is respectively 0.3803 and 0.2244. The difference in the FCI between the two species is highly significant (two-sided t-test p=9.796e-12). Within rat pyramidal neurons, layer 5 pyramidal neurons are significantly more complex than layer 2/3 pyramidal neurons (Figure 2B, green, two-sided t-test p=0.048). Interestingly, this is not the case in humans, where layer 2/3 pyramidal neurons are significantly more complex both compared to layer 4 (two-sided t-test p=0.013) and layer 5 (two-sided t-test p=0.010) pyramidal neurons (Figure 2B, orange). It is interesting to note that in the human cortex, layer 2/3 is expanded relative to layer 5 (Galakhova et al., 2022), and contains several novel cell types (Berg et al., 2021 and see **Discussion**).

**Figure 2.**
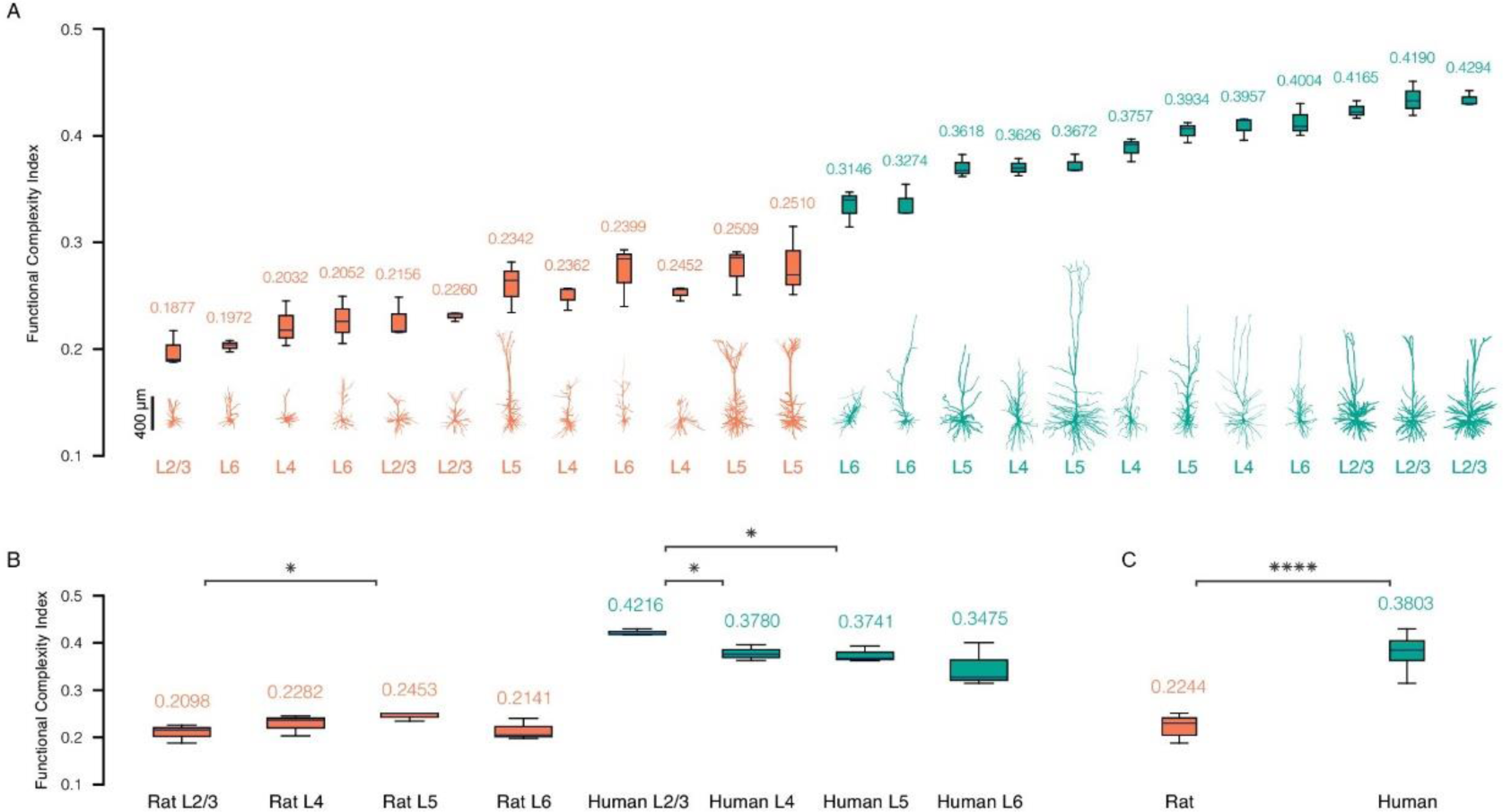
Human cortical pyramidal neurons are more functionally complex compared to rat cortical pyramidal neurons. **A.** Top: Functional complexity index (FCI) scores for all 24 (12 rat in orange and 12 human in green) modeled neurons depicted alongside with their respective morphology (bottom). **B.** Comparison of FCI per cortical layer for rat neurons (orange) and human neurons (green). **C.** Overall comparison of the FCI between the two species, see **Methods** and Table S1 for morphological details. * p value smaller than 0.05, and **** p value smaller than 0.0001.

In summary, Figure 2 demonstrates that our new index for assessing the functional complexity of neurons is sensitive enough to capture variation between cortical pyramidal neurons across cortical layers and across species.

What are the specific factors that contribute to the greater functional complexity of human neurons? To answer this question, we first examined whether morphological properties *per se* are responsible for the greater complexity of human cortical pyramidal neurons. To that end, we repeat our FCI assessment process, only now assigning rat type synapses to all morphologies, both human and rat (see **Methods**). Compared to Figure 2 where we used rat synapses for rat models and human synapses for human models, here the difference between species is less pronounced, although human neurons still exhibit statistically significant higher FCI, on average (Figure S5, two-sided t-test p=0.022). This implies that some morphological features contribute to the extra-complexity of human neurons, in addition to the crucial role of synaptic properties that we examine later in Figure 4.

We next extracted 58 different morphological features for the modeled neurons (see **Methods**). In particular, we characterized morphological features related to *trunk branches* (branches that emerge from the soma and end in a bifurcation) as well as features related to *termination branches* (branches starting from a bifurcation and ending at the dendritic tip), and *bifurcation branches* (all other branches – those that start and end in a bifurcation), see Figure 3A for a graphical demonstration. In order to study which morphological features best predict the FCI, we computed the correlation between the FCI and each of the 58 features measured (Figure 3B-E, I). Figure 3I presents a histogram of the *R*^2^ correlation values between the FCI and individual features. It is evident that only a few specific features explain a substantial portion of the FCI’s variance. The single feature best predicting the FCI was the entire area of the dendritic tree (*total dendritic area*), with *R*^2^ = 0.74 (Figure 3B). The total length of bifurcating branches (orange branches in Figure 3A) achieved an *R*^2^ = 0.45 (Figure 3C), whereas *longest bifurcation branch*, which is closely related to the maximal path distance of the tree from soma to tip, achieved an *R*^2^ = 0.44 (Figure 3D). Surprisingly, the feature reflecting the number of bifurcation branches, achieved a modest *R*^2^ of 0.29 (Figure 3E).

**Figure 3.**
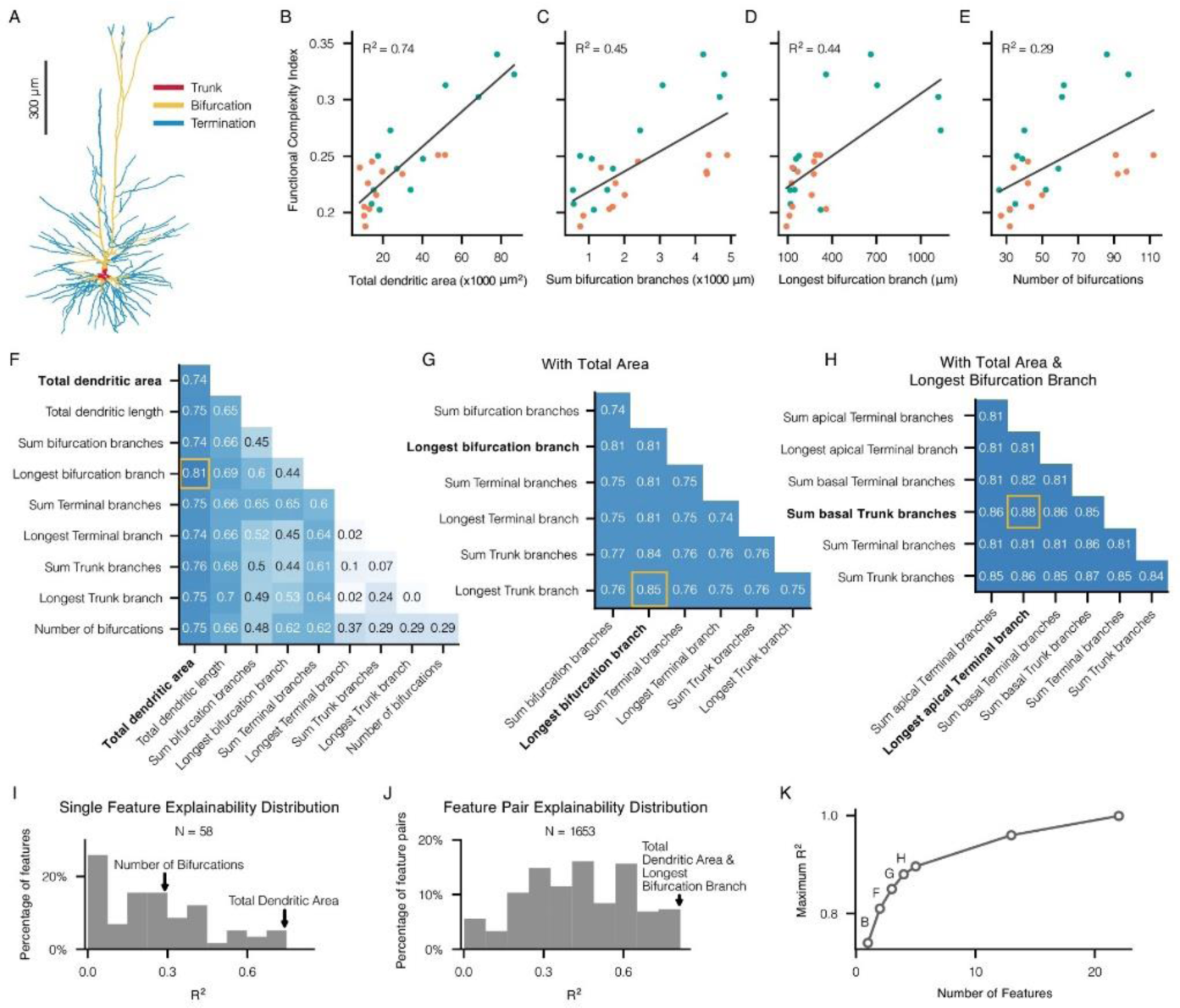
Correspondence between morphological features and the functional complexity index. **A.** Human layer 2/3 dendritic tree colored by three dendritic subtrees (trunk, bifurcation and termination) as indicated at top left. **B-E.** Correlation between single morphological features and FCI; green circles for human neurons and orange circles for rat neurons. **F.** Correlations between FCI with pairs of morphological features (the diagonal refers to the correlation with the single feature). Yellow square highlights the largest correlation. **G.** Correlations between FCI and triplets of morphological features. Each pair depicted already incorporates the total dendritic area. **H.** FCI correlation with quadruplets of morphological features; each case includes the total dendritic area and the longest bifurcation branch. **I.** Distribution of FCI correlation with single morphological features. **J.** Distribution of FCI correlation with a pair of morphological features. **K.** Maximal correlation achieved using increasing numbers of morphological features; the cases corresponding to B, F, G and H are marked above the graph.

Next, we asked what is the minimum number of combined features that most closely predict the FCI. Figure 3J displays a histogram of the *R*^2^ values showing how well pairs of features account for the complexity index. This analysis reveals that pairs of features, when considered together, generally provide a greater explanation of the complexity variance than individual features. Figure 3F illustrates the *R*^2^ values explained by different feature pairs. Notably, the most predictive pairs consistently included *total dendritic area*. The most predictive pair also included *longest bifurcation branch*, that together with *total dendritic area* achieved *R*^2^ of 0.81. In Figure 3G, we correlated the FCI with triplets of features, each including *total dendritic area*. Again, all triplets best predicting the FCI always included *longest bifurcation branch*, with the best triplet attaining a value of *R*^2^ = 0.85 (*total dendritic area, longest bifurcation branch* and *longest trunk branch*). Finally, in Figure 3H, we correlated the FCI with quadruples of features, containing both *total dendritic area* and *longest bifurcation branch*. The best predicting quadruple achieved a value of *R*^2^ = 0.88. Overall, the best third and fourth features were related to either apical or basal trunk branches and terminal branches. Importantly, the coefficients of the third and fourth features were negative, suggesting that the less dendritic length is invested in trunk branches and terminal branches, the more complex the neuron is. In other words, the greater the dendritic length allocated to bifurcation branches, the more complex the neuron becomes (see **Discussion**). Using additional features beyond this core group of four features marginally contributes to the variance explained (Figure 3K). Notably, the full complexity can be entirely explained using a total of 23 features (Figure 3K, top point at right).

In Figure 4, we explore the impact of synaptic properties that, as mentioned above, contribute to the increased complexity of human pyramidal cells compared to rat pyramidal cells. To that end, we repeated our FCI assessment process, only now assigning each morphology with either one of four different synaptic types: rat synapses, human synapses, and two hybrid variants that attempt to disentangle the specific contributions of synaptic conductance of AMPA + NMDA channels, and NMDA γ factor values that are related to the steepness of the NMDA nonlinearity (see Equation (6) in **Methods**). The hybrid A synaptic type has the rat type synapse parameters (including rat conductance values) together with the human γ factor, whereas the hybrid B synaptic type has the human type synapse parameters (including human conductance values), together with the rat γ factor (see **Methods** and Table S2).

**Figure 4.**
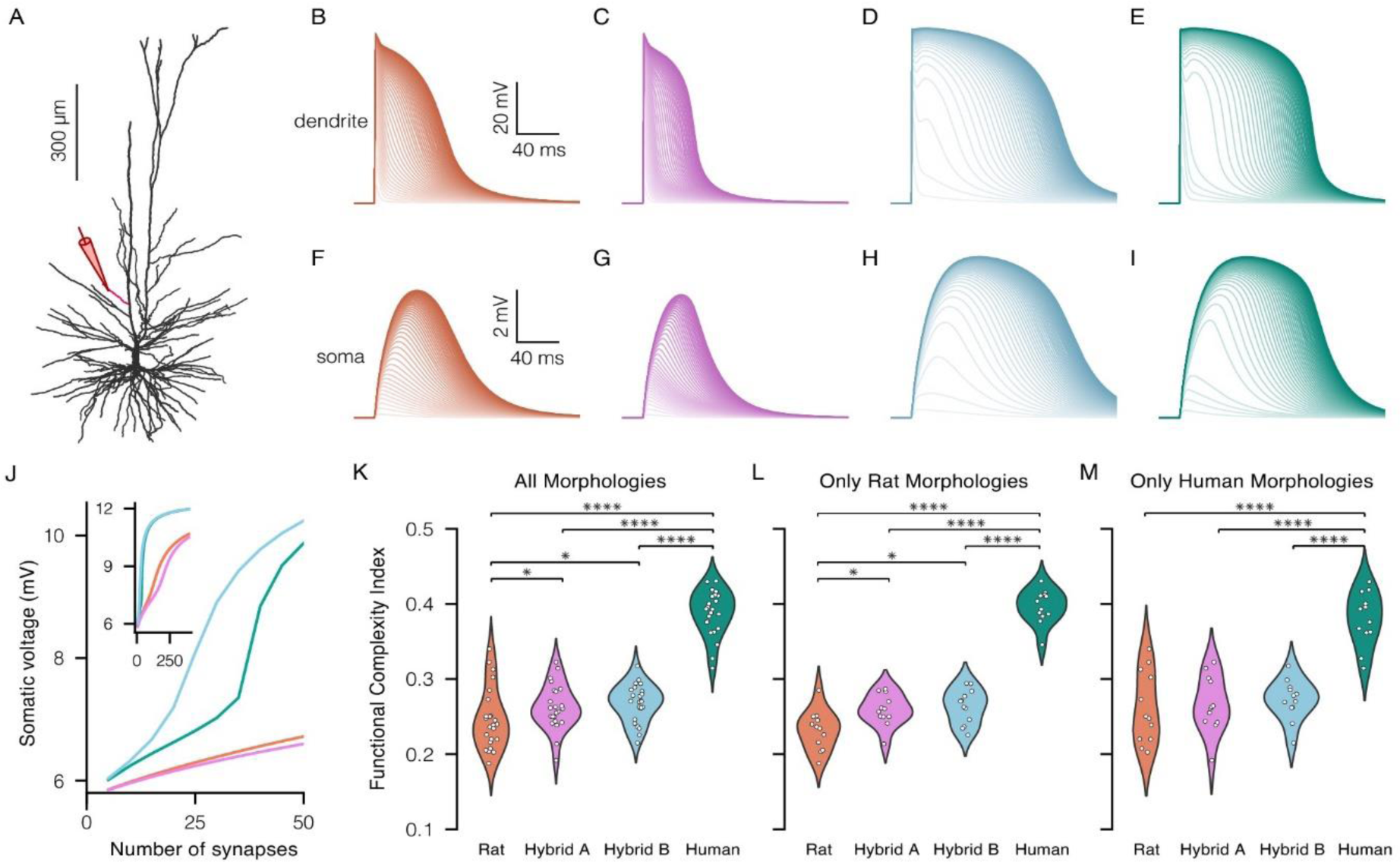
Correspondence between various synaptic features and the functional complexity index. **A**. Modeled human layer 2/3 pyramidal neuron; the oblique branch receiving excitatory synapses is depicted in red with yellow electrode at left. **B-E** Local dendritic voltage responses in the activated oblique branch of the modeled cells shown in A for different synaptic types; when increasing the numbers of simultaneously activated synapses (from 5 synapses to 400 synapses) **B**. Rat synapses where used (orange). **C**. Rat synapses with the γ factor of human for the NMDA conductance (pink). **D**. Human synapses with γ factor of rat (blue). **E.** Human synapses were used (green). **F-I** as in B-E but the respective soma voltage response. **J.** Somatic EPSP amplitude as a function of the number of activated dendritic synapses, for the four cases shown in F-I. **K.** FCI distribution for all (human and rat) 24 morphologies, for the 4 different synapse types/cases (colours as in B-I). **L.** FCI distribution for rat morphologies, with the different synapse types. **M** FCI distribution for human morphologies, given different synapse types, see **Methods**). * p value smaller than 0.05, and **** p value smaller than 0.0001.

In order to investigate the NMDA receptor nonlinearity across various synaptic types, we progressively activated an increasing number of synapses along a dendritic segment of a representative human layer 2/3 neuron model (Figure 4A), examining the four distinct synaptic types. The resulting local dendritic responses are illustrated in Figure 4B-E, whereas the corresponding somatic voltage responses are depicted in Figure 4F-I. Figure 4J shows the peak somatic voltage as a function of the number of activated synapses on a single oblique branch of a human L2/3 model as shown in Figure 4A, for each of the 4 synapse types used. The rat and hybrid A type synapses exhibited linear responses with fewer than 50 simultaneous synaptic activations. The inset demonstrates that in these two cases, the NMDA response is saturated with this number of activated synapses. However, with the same number of 50 activated synapses, both human and hybrid B type synapses demonstrated a significant increase in somatic voltage response, which results from the generation of highly nonlinear NMDA spike in the activated oblique dendrite. Notably, the human synapse type exhibited a critical transition to steep nonlinearity around the activation of 35 synapses, shifting from sublinear to supralinear summation of synaptic inputs.

Indeed, we found that models with human type synapses were significantly more complex than models with rat type synapses, across rat morphologies (Figure 4L), across human morphologies (Figure 4M) and across all morphologies together (Figure 4K). However, models with either hybrid A or hybrid B synaptic types were only slightly more complex than models with rat synaptic types. These findings are consistent with the results shown in Figure 4J, highlighting the impact of the more nonlinear NMDA receptors on the complexity of human neurons.

We conclude that the contribution of synaptic properties to the increased complexity of human pyramidal cells, compared to rat, is primarily driven by the enhanced nonlinearity of the NMDA receptor dynamics.

## Discussion

Human neurons exhibit distinct structural and biophysical properties compared to those of rats, yet it is unclear whether these differences translate into a greater functional complexity at the system level that could explain humans’ elevated cognitive abilities. Utilizing a deep learning-based framework, we developed a novel generalized Functional Complexity Index (FCI) to systematically assess the input-output complexity of neurons. Using the FCI, we demonstrated that human cortical pyramidal neurons are significantly more functionally complex than their rat counterparts, suggesting a link between neuronal complexity and enhanced cognitive abilities in humans. This is due to differences in both morphological features and dynamics of excitatory synapses. In particular, we have shown that human neurons are functionally more complex thanks to their larger surface dendritic area and extensive bifurcation patterns. In addition, human NMDA-activated receptors exhibit steeper nonlinear voltage responses, enabling more complex I/O relationship.

Since the seminal studies of W. Rall (1959, 1964, 1977), highly realistic fine-scale biophysical models of individual neurons were constructed across brain regions and across species (Hay et al., 2011; Markram et al., 2015; Allen Institute for Brain Science, 2015, Eyal et al., 2018, Hunt et al., 2023). These models reflected the remarkable variability in morphology and physiology of cortical neurons, both within and between species, human cortex included. Despite significant progress in “understanding neurons”, a crucial gap persisted: we lacked a systematic, quantitative tool to measure the functional complexity of neurons; namely the complexity of their I/O relationship. Such measure is key for comparing neurons’ complexity across different neuronal types, layers and species, but most crucially, for connecting the computational complexity of single neurons to that of the neuronal network.

Our proposed FCI measure addresses this issue by quantifying the functional complexity of the I/O function of neurons at the synapse(input)-to-spike(output) resolution, based directly on their morpho-electrical properties. Unlike previous methods, such as Poirazi and Mel (2001), that used abstract models to infer memory capacity, our approach directly evaluates the complexity of detailed biophysical models of neurons. Since these models more closely match real neurons, the FCI provides a biologically accurate representation of their computational capacity. Compared to compartmental modeling studies, like Eyal et al. (2018), that focused on the number of nonlinear dendritic subunits, the FCI provides greater scalability by using deep learning techniques (Beniaguev et al., 2021) to generalize across various neuron types and species. Additionally, whereas previous work limited its scope to subthreshold dynamics (e.g., Ujfalussy et al. 2018), the FCI captures both suprathreshold and subthreshold behaviors.

It is worth mentioning that rather than using the depth of the analogue DNN of the respective biophysical neuron model as a proxy for its complexity, the FCI is evaluated based on the accuracy of a fixed-size DNN (three layers in this study) in matching the I/O of the biophysical model. This provides a more precise and interpretable metric for assessing the functional complexity of neurons and understanding the underpinning of this complexity. A neuron that achieves a larger FCI score typically requires a deeper DNN to accurately replicate its I/O behavior, thus maintaining the connection between complexity and network depth. Consequently, the FCI can be viewed as a measure of how “deep” a neuron’s computational capabilities are, analogue to how deeper artificial neural networks capture more complex patterns. In this sense, a higher FCI value reflects the neuron’s capability of performing more complex, layered processing.

The fixed DNN architecture used to assess neuron complexity makes the FCI a robust, interpretable measure. It enables a systematic comparison of neurons, revealing how their morphology and biophysics shape their functional complexity. However, this measure faces several challenges, notably the computational cost of generating large I/O datasets from the biophysical model and training the respective neural networks, especially when varying biophysical parameters and morphology of neurons. Additionally, the use of output normalization in the FCI (see **Methods**) focuses the sampling on a particular regime of the model’s I/O space. Moreover, the method depends on specific hyperparameters and DNN architecture, which might influence accuracy and introduce variability in the value of the respective FCI. Overly expressive architectures may reduce complexity differences, whereas under-expressive ones may inflate them, misrepresenting simpler neurons. Careful architecture selection is crucial to avoid overfitting or oversimplification and to ensure a meaningful dynamic range. Notice that the numerical values of the FCI depend on the specific DNN architectural choices, making it a measure that is relative to the selected architecture.

To address these issues, we used a three-layer temporally convolutional network (TCN), a DNN architecture that has been shown to successfully predict the I/O function of a biophysically detailed model of rat L5 pyramidal cell across multiple scenarios (Beniaguev et al., 2021). Furthermore, we validated the robustness of our approach by testing a subset of neurons with slightly different architectures, a two-layer TCN and a seven-layer TCN instead of a three-layer TCN. We found that the rankings of neuron complexities remained consistent with the ranking presented in our results (not shown). This demonstrates that the method reliably captures complexity differences across neuronal types and species.

We found that the increase in FCI in humans is correlated with a larger surface area of the dendritic tree, larger dendritic tree height (soma-to-tip distance), and a greater proportion of the dendritic length allocated to bifurcation branches (Figure 3). The larger dendritic tree size combined with the increased allocation of dendritic length to bifurcation branches possibly enables greater compartmentalization, allowing distinct regions of the dendritic tree to process inputs semi-independently, enhancing computational capacity (Polsky et al., 2004; Beualieu-Laroche et al., 2018; Eyal et al., 2018; Beualieu-Laroche et al., 2021; Otor et al., 2022). It is worth noting that without considering the features related to tree size, the correlation of the number of branches *per se* is rather poor. While previous works (Poirazi and Mel, 2001) emphasized the number of independent subunits as a key factor for memory capacity, our results suggest an interaction between tree size and bifurcation pattern that determines the number and the level of independence of subunits.

Human neurons have larger dendritic spine head area compared to rodents (Benavides-Piccione et al., 2002; Ofer et al., 2022) and correspondingly, more NMDA receptors per synapse (Eyal et al., 2018; Hunt et al., 2023) with stronger nonlinear voltage-dependent dynamics (Eyal et al., 2018; but see Testa-Silva et al., 2022). These synaptic properties enable stronger and larger combinatorial interactions between local excitatory synapses; this contributes to a more nonlinear and complex I/O relationship and thus to a larger FCI (Figure 4). These findings agree with previous research linking synaptic nonlinearity to functional complexity (Mel, 1992; Mel, 1994; Larkum et al., 1999; Schiller et al., 2000; Branco et al., 2010; Major et al., 2013; Larkum et al., 2020).

These morpho-biophysical features contributing to neuronal complexity are also reflected in differences in FCI value across cortical layers. Human layer 2/3 pyramidal neurons exhibit greater complexity than neurons in other layers, including the large layer 5 neurons (Figure 2). This is an opposite pattern to that observed in rats, where layer 5 pyramidal neurons are the most complex. It was shown that human cortical layer 2/3 is expanded relative to other cortical layers, including layer 5 (Galakhova et al., 2022). Taken together, these findings suggest that humans have more layer 2/3 neurons, each of which is individually more complex. This might relate to the increased, and potentially novel (Berg et al., 2021), role of layer 2/3 in human cortical computation.

Future research could expand this study to explore the impact of active dendritic properties, such as those of voltage-dependent Na^+^ and Ca^+2^ ion channels on the FCI, as these channels have unique properties in human dendrites (Gidon et al., 2020; Gooch et al., 2022). Unfortunately, accurate models with dendritic nonlinear conductance validated against experimental data remain quite rare, highlighting the need for further advancements in this area. Also warrants further investigation is the impact of the abundant dendritic spines on the I/O transformation of human cortical neurons (Yuste et al., 1995; Benavides-Piccione et al., 2002; Elston et al., 2003), their unique axonal excitability (Wilbers et al., 2023) and local connectivity patterns (DeFelipe, 2011; Oh et al., 2014; Loomba et al., 2022; Shapson-Coe et al., 2024). Another worthy direction is to extend this study onto additional neuronal types such as hippocampal CA1 and CA3 pyramidal neurons and cerebellar Purkinje cells. Studying the FCI in neurons of other species (e.g., non-human primates) and exploring how the functional complexity of neurons (the FCI) impact network-level computations would deepen our understanding of how neuronal diversity impact cognitive capabilities.

## Methods

### Neuron morphologies

Morphologies of 24 3D-reconstructed cortical pyramidal neurons were used in this study, 12 rat pyramidal cells and 12 human pyramidal cells. 3 neurons were modeled from each of the following layers: layer 2/3, layer 4, layer 5, layer 6. Rat neurons were taken from (Hay et al., 2011; Markram et al., 2015; Reimann et al., 2024) and human neurons from (Mohan et al., 2015; Allen Institute for Brain Science, 2015). To consider the variability in the reconstruction quality, the diameters of all morphologies were edited such that no diameter would be smaller than 0.3 μ*m*. A complete description of the morphologies used is provided in Supplementary Table 1.

### Neuron models

We constructed a detailed biophysical model (Rall, 1964) for each morphology. All models have specific membrane capacitance Cm = 1μF/cm^2^, specific axial resistance *R*_*a*_ = 150 Ω *cm* and specific membrane resistance *R*_*m*_ = 20,000 Ω cm^2^. All models were equipped with spike-generating voltage-dependent Na^+^ and K^+^ ion channels in the soma and axon. Channel kinetic is as in Hay et at. (2011). The maximal conductance of the active channels of all models was fit to match the experimental F-I curve as in Hay et al. (2011). The maximal conductance of the active channels in the soma and axon of all morphologies were normalized by the electrical load that the dendritic tree imposes on the soma (ρ_*soma*_) and on the axon (ρ_*axon*_), using the rho scaling method (Hay et al., 2013). By this, the conductance of each somatic or axonal active channel for each morphology was set as follows:

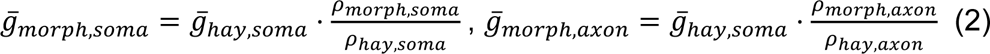

Where ρ is the dendrite-to-soma or dendrite-to-axon conductance ratio defined as:

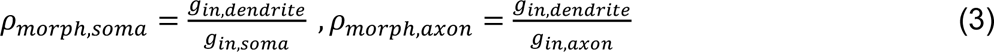

### Synapse models

For each neuron model, one excitatory AMPA + NMDA-based synapse and one inhibitory GABA_A_-based synapse were placed on every 1μ*m* dendritic length. The synaptic current was modeled as:

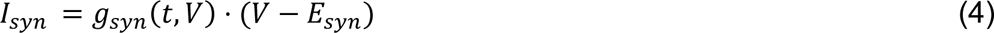

Where *E*_*syn*_ is the reversal potential for the synaptic current and *g*_*syn*_ is the synaptic conductance modeled using two-state kinetic scheme:

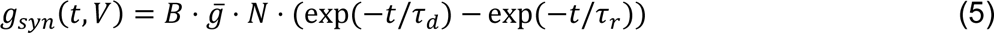

Here 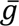 is the peak conductance and *N* is a normalization factor given by:

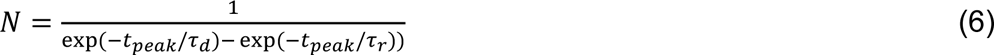

where *t*_*peak*_, time to peak of the conductance, is:

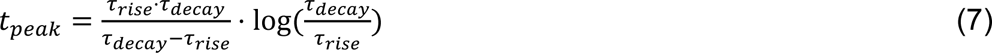

Where τ_rise_ and τ_decay_ are the rise time and decay time constants. For AMPA and GABA_A_ conductances, *B* = 1 (voltage-independent conductance).

For the voltage-dependent NMDA conductance B was defined as in Jahr and Stevens (1990):

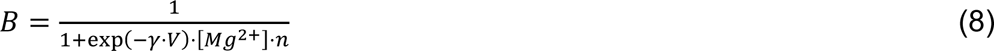

[*Mg*^2+^] was set to 1 *mM*, n was 1/3.57mM. The kinetics (synaptic rise and decay time constants, etc.) and conductances of rat synapses were taken from Markram et al. (2015), while those of human synapses were taken from Eyal et al. (2018). The “hybrid A” and “hybrid B” type synapses included a mix of rat and human synaptic properties. A full description of the synaptic properties is provided in Supplementary Table 2.

### Normalizing for the input firing rates

In order to avoid the possible confounding effect of the different firing rates of different models on the FCI, we carefully selected the rate of the input excitatory (E) as well as inhibitory (I) synapses such that the average output firing rate of all models will be 1 sp/s. For each model, we chose 10 valid input E/I firing rate combinations that resulted in an average output firing rate that is within 0.01 sp/s around the chosen 1 sp/s mark. Every valid input firing rate combination spans a range of 0.1 sp/s difference in firing rate both in excitation and in inhibition (for example, a valid input firing rate combination of a specific model might be 1-1.1 sp/s in excitation and 2-2.1 sp/s in inhibition, which amounts for an average output firing rate of 1.005 sp/s). To find valid input firing rate combinations, we exhaustively searched the input firing rate space between 0 sp/s to 20 sp/s in both excitation and inhibition (Figure S3).

### Simulations and resulting datasets

In order to fit DNN models per simulated neuron, we followed the study of Beniaguev et al. (2021). First, we generated a simulation dataset for each modeled neuron. In each simulation, the modeled neuron was stimulated by random excitatory and inhibitory synaptic input (one synapse per 1μ*m* dendritic length) distributed randomly over the dendritic surface of the modeled neuron for a duration of 10 s. As explained above, in each simulation we used an input regime that results in an output firing rate of ∼1 sp/s. Each presynaptic spike train was sampled from a Poisson process with a smoothed piecewise constant instantaneous firing rate. The Gaussian smoothing sigma, as well as the time window of constant rate before smoothing, were independently resampled for each 10 s simulation from the range of 10 ms to 1000 ms. This was the case, as opposed to choosing a constant firing rate, to create additional temporal variations in the data, in order to increase the applicability of the results to a wide range of potential input regimes. For each neuron model, we created a dataset consisting of 12,000 train simulations of 10 s each, equivalent to ∼1.4 days of neural data (see below). Simulations were performed using NEURON software (Carnevale and Hines, 2006) and were run in parallel on a CPU cluster.

### Fitting I/O of neuron models to respective DNNs

We followed Beniaguev et al. (2021) to train DNNs based on the neuron model simulation datasets. The DNN was fed as an input with the same presynaptic spike as the biophysical model did. The respective DNN was expected to produce voltage output that matches as closely as possible both the subthreshold and the spiking activity at the soma. In this study, we predefined a fixed-size temporally convolutional network (TCN) with 3 layers and a width of 128 units per layer for all neuron models (Beniaguev et al., 2021; Bai et al., 2018) with 3 different random initializations per modeled neuron. For a subset of the neuron models, we also fit two-layer TCNs, and repeated our measurements as explained above. We found that selecting different TCNs as a benchmark, did not affect the results qualitatively. Namely, the ranking of the neuron’s complexity remained almost the same when changing the depth of the respective TCN. Each network was trained for approximately 4 days of neural data, corresponding to roughly 3 full epochs over the entire training dataset. The total number of single GPU years needed to fit all DNNs throughout the entire study was ∼2.3 years.

### DNN performance

We divided our 12,000 simulations to a training set of 10,000 simulations, a validation set of 1,000 simulations and a test set of 1,000 simulations. We fitted all DNN models on the training set and calculated the DNN performance on the unseen test set. The validation set was used for modeling decisions, hyperparameter tuning and snapshot selection during the training process (early stopping). The DNN’s task was the binary classification task of predicting whether the neuron emitted a spike in all 1 ms time points. This was evaluated using the receiver operator characteristic (ROC) of binary spike prediction. The performance was finally quantified using the area under the curve (AUC) of the ROC. Additional details are found in Beniaguev et al. (2021).

### Functional Complexity Index

We defined the Functional Complexity Index (FCI) of a neuron model as inversely proportional to the performance of its respective DNN (Figure 1 and Equation (1)). Specifically, the performance of the DNN model was quantified using the Area Under Curve (AUC) measure. We found that typical values of AUC of such models ranged between 0.9 to 0.999 (Beniaguev et al., 2021). In other words, an *AUC* = 0.9 indicates a very poor performance of the DNN. Therefore, the *FCI* of such cases was set to 1 (Equation (1)). For a great performance where the *AUC* = 0.999, the FCI was set to 0 (see Figure S4 for the full relationship between the FCI and the AUC).

### Morphological features

We used NeuroM (Arnaudon et al., 2024) to calculate the values of various morphological features for each of our modeled morphologies. The following features were considered: total dendritic length, total dendritic area, number of forking points, number of bifurcation points (a forking point of exactly two branches), number of leaves, max Radial distance, max branch order, mean sibling ratio, sum/mean/longest bifurcation branches, sum/mean/longest terminal branches and sum/mean/longest trunk branches. Each of these features was calculated separately for the basal and the apical trees of each morphology. Additionally, we used three features related to the entropy of the topological representation of the dendritic tree (Kanari et al., 2018), namely, the sum/mean/max entropy of the morphology. In total, we had 58 morphological features.

### Correlation between morphological features and complexity

To predict the value of the FCI from the neuron’s morphological features, we used linear regression to fit the following equation:

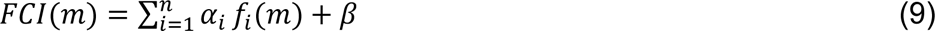

where *f*_*i*_(*m*) is the *i*-th feature computed for a given morphology, *m*. *α*_*i*_ is the fitted coefficient for the *i*-th feature; *β* is a fitting bias and *n* is the number of features used for fitting. In this study, we computed Equation (7) with *n* ranging from 1 to 4.

Given a linear regression curve, we calculate the *R*^2^ to quantify how well this curve fits the data. The results for different numbers of features (*n*) are provided in Figure 3. In Figure 3F-H, yellow square indicates the highest correlation.

## Acknowledgments

We thank Oren Amsalem for his early work on the functional complexity index. We thank all lab members of the Segev and London Labs for many fruitful discussions and valuable feedback regarding this work. This work was supported by the ONR grant award number N00014-24-1-2055 and grant award number N00014-23-1-2051. M.L. was supported by the ISF grant 1331/23, the NIPI grant 206-22-23, and the BSF grant 2023104. I.S. was supported by the Drahi Family Foundation, the ETH domain for the Blue Brain Project, the Gatsby Charitable Foundation and the NIH grant agreement 1RM1NS132981-01.

## Author contributions

I.A., conceptualization, methodology, investigation, visualization, software, validation, data curation, writing – original draft; D.Y., investigation, visualization, software, writing – original draft; D.B., conceptualization, methodology, writing – review & editing; C.P.J.K., methodology, investigation, validation, data curation, writing; I.S. and M.L., conceptualization, methodology, writing – review & editing, supervision, resources, funding acquisition.

## Competing Interests statement

The authors declare no competing interests.

## Code availability

The simulation, fitting, and FCI calculation code are publicly available on GitHub (http://github.com/ido4848/fci).

## Data availability

The neuron morphologies and neuron models appearing in Figure 1 (Rat L2/3 and Human L2/3), as well as two additional morphologies and models (Rat L5 and Human L5) are publicly available on GitHub (http://github.com/ido4848/fci). All other neuron morphologies and neuron models are available upon request. All spike times and somatic membrane potentials presented in the article are available upon request. All FCI values and correlation values presented in the article are available upon request.

---

**Supplementary Figure 3.**
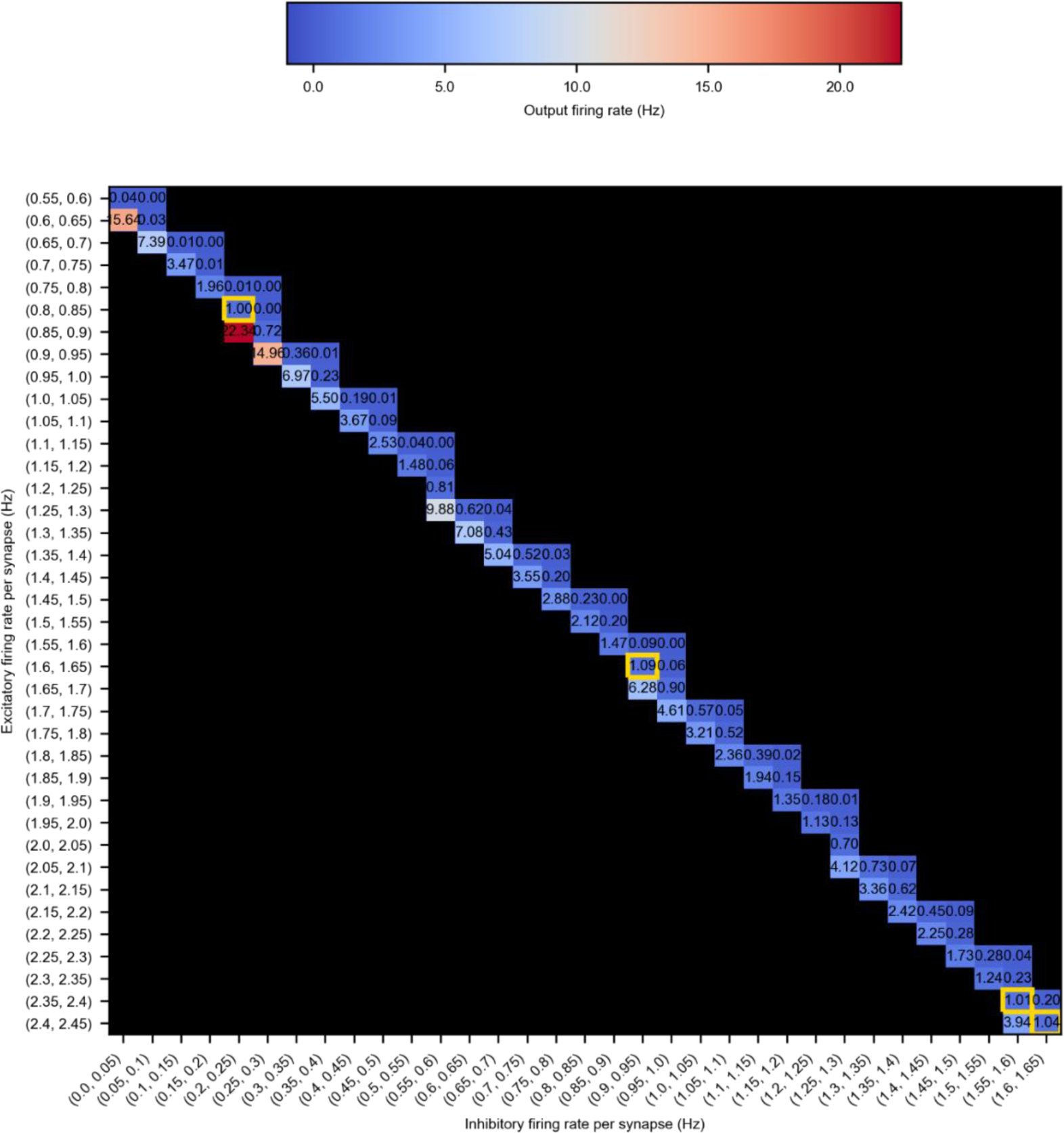
Io matrix.

**Supplementary Figure 4.**
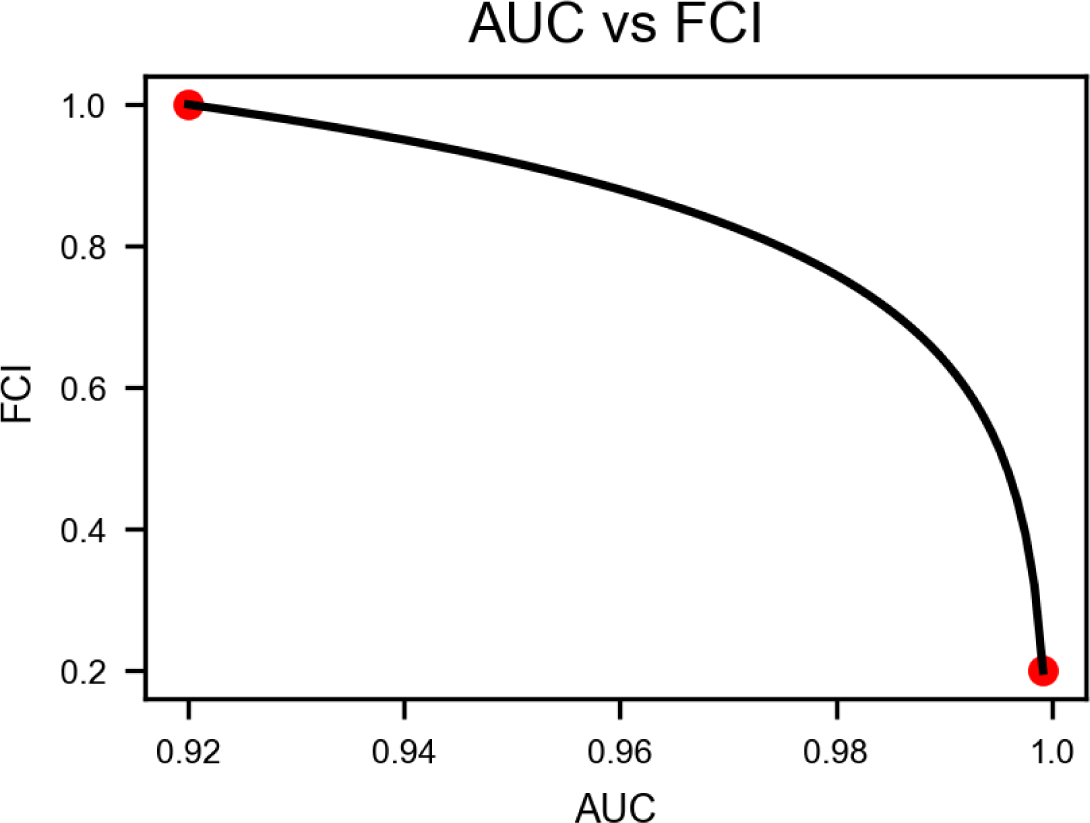
Relation between FCI and AUC.

**Supplementary Figure 5.**
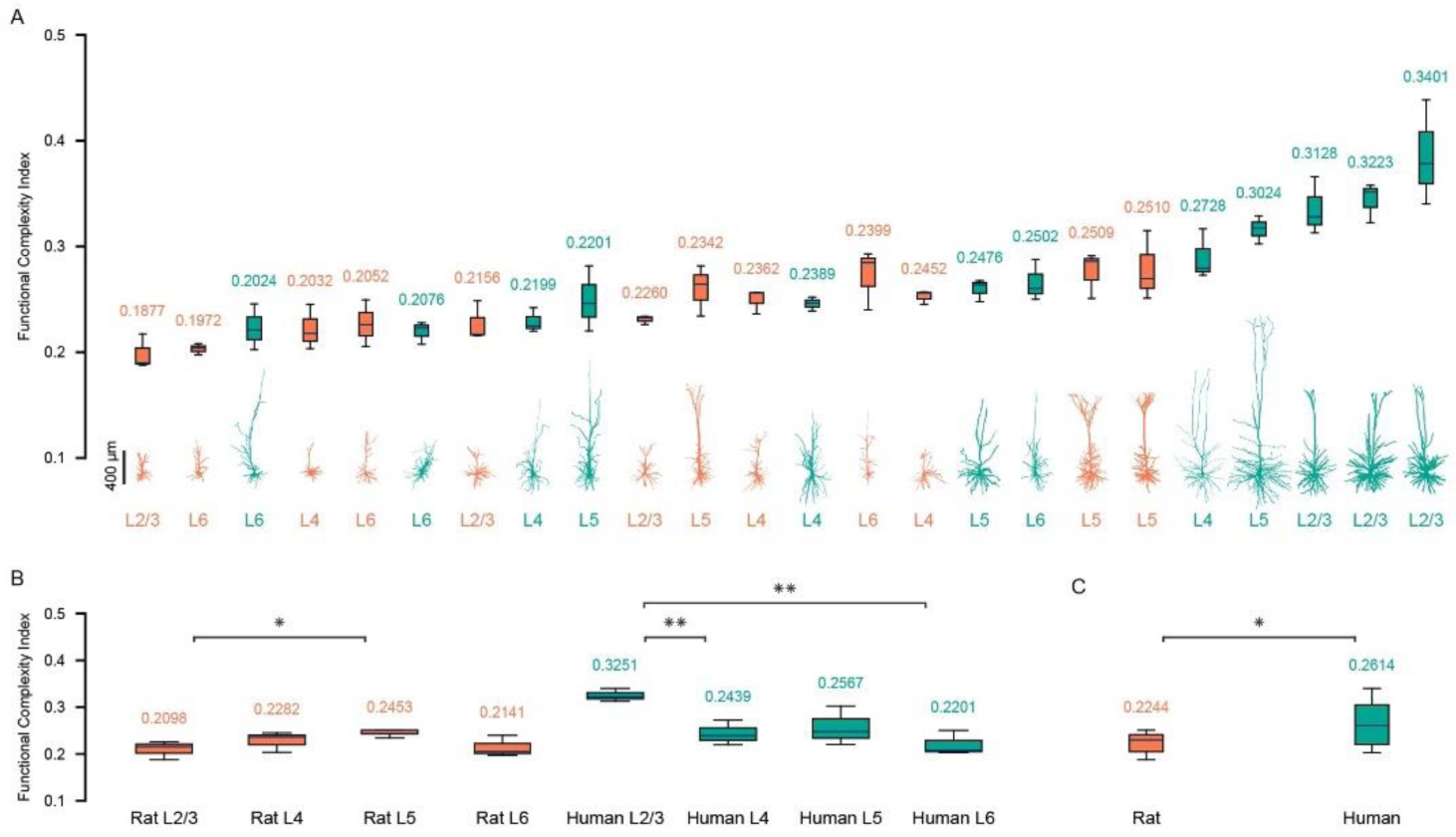
FCI of all morphologies with rat synapses.

**Supplementary Table 1.** morphologies.

| # order in Figure 2 | Species | Cortical layer | Morphology identifier | Citation |
| --- | --- | --- | --- | --- |
| 1 | Rat | L2/3 | L2 TPC | Reimann et al., 2024 |
| 2 | Rat | L6 | L6 IPC | Reimann et al., 2024 |
| 3 | Rat | L4 | L4 TPC | Reimann et al., 2024 |
| 4 | Rat | L6 | L6 TPC | Reimann et al., 2024 |
| 5 | Rat | L2/3 | 229_5 | Markram et al., 2015 |
| 6 | Rat | L2/3 | 229_1 | Markram et al., 2015 |
| 7 | Rat | L5 | cell1 | Hay et al., 2011 |
| 8 | Rat | L4 | 230_1 | Markram et al., 2015 |
| 9 | Rat | L6 | L6 UPC | Reimann et al., 2024 |
| 10 | Rat | L4 | 230_2 | Markram et al., 2015 |
| 11 | Rat | L5 | TTPC_1 232_1 | Markram et al., 2015 |
| 12 | Rat | L5 | L5 TPC | Reimann et al., 2024 |
| 13 | Human | L6 | 548494556 | Allen Institute for Brain Science, 2015 |
| 14 | Human | L6 | 528614014 | Allen Institute for Brain Science, 2015 |
| 15 | Human | L5 | 1833 | Mohan et al., 2015 |
| 16 | Human | L4 | 539661667 | Allen Institute for Brain Science, 2015 |
| 17 | Human | L5 | 2057 | Mohan et al., 2015 |
| 18 | Human | L4 | 569818704 | Allen Institute for Brain Science, 2015 |
| 19 | Human | L5 | 790872626 | Allen Institute for Brain Science, 2015 |
| 20 | Human | L4 | 1496 | Mohan et al., 2015 |
| 21 | Human | L6 | 558211203 | Allen Institute for Brain Science, 2015 |
| 22 | Human | L2/3 | 1204 | Mohan et al., 2015 |
| 23 | Human | L2/3 | 1148 | Mohan et al., 2015 |
| 24 | Human | L2/3 | 1125 | Mohan et al., 2015 |

**Supplementary Table 2.** synaptic parameters.

| synapse type | AMPA |  |  | NMDA |  |  |  | GABA A |  |  |
| --- | --- | --- | --- | --- | --- | --- | --- | --- | --- | --- |
| parameter | tau_r | tau_d | g_max | tau_r | tau_d | gamma | g_max | tau_r | tau_d | g_max |
| units | ms | ms | nS | ms | ms | 1/mV | nS | ms | ms | nS |
| rat | 0.2 | 1.7 | 0.4 | 0.29 | 43 | 0.062 | 0.3 | 0.2 | 8 | 0.7 |
| human | 0.3 | 1.8 | 0.88 | 5 | 43 | 0.078 | 1.31 | 0.2 | 8 | 0.7 |
| hybrid A | 0.2 | 1.7 | 0.4 | 0.29 | 43 | 0.078 | 0.3 | 0.2 | 8 | 0.7 |
| hybrid B | 0.3 | 1.8 | 0.88 | 5 | 43 | 0.062 | 1.31 | 0.2 | 8 | 0.7 |

## Notes

https://github.com/ido4848/FCI

### Competing Interest Statement

The authors have declared no competing interest.

### Summary of Updates

There was an error in the PDF that is possibly fixed. Removed line numbers.

